# Circadian regulation of hippocampal function is disrupted with chronic corticosteroid treatment

**DOI:** 10.1101/2022.07.12.499774

**Authors:** Matthew T. Birnie, Matthew D.B. Claydon, Benjamin P. Flynn, Mitsuhiro Yoshimura, Yvonne M. Kershaw, Rebecca C.R. Demski-Allen, Gareth R.I. Barker, E. Clea Warburton, Zuner A. Bortolotto, Stafford L. Lightman, Becky L. Conway-Campbell

**Author notes:** Correspondence to: Matthew T. Birnie, University of Bristol, Henry Wellcome Laboratories for Integrative Neuroendocrinology, Whitson St, Bristol, BS1 3NY. These authors contributed equally to this work (Matthew T. Birnie, Matthew D.B. Claydon).

## Abstract

Neuropsychiatric disorders such as major depression and schizophrenia are highly prevalent and contribute substantially to disease burden worldwide. Despite this, progress understanding the pathophysiology has remained largely elusive, yet these disorders often exhibit a loss of regulation of biological rhythms, such as sleep/wake cycles and hormonal rhythms. Cushing’s disease, a condition characterized by chronic corticosteroid (cortisol) hypersecretion is associated with psychiatric and neurocognitive disorders and disruption to the circadian release of cortisol can result in depression and neurocognitive impairment. In rats, we report that circadian regulation of the hippocampal transcriptome integrates crucial functional networks that link corticosteroid-inducible gene regulation to synaptic plasticity regulation via an intra-hippocampal circadian transcriptional clock. During the early active period, when corticosteroid availability is high, CA1 region excitatory and inhibitory post-synaptic currents were augmented along with long-term potentiation. In contrast, chronic corticosteroid exposure disturbed hippocampal function. The hippocampal transcriptome, as well as circadian regulation of synaptic plasticity were ablated, resulting in memory loss during hippocampal-dependent behavior. These findings identify how exposure to elevated levels of corticosteroid, that is often seen in neuropsychiatric illness, results in adverse critical hippocampal function. These data provide novel insights into the molecular mechanisms of neurocognitive disorders and provides evidence for corticosteroid-mediated intervention in disabling mental illnesses.

## Main

Circadian rhythms regulate many physiological, biological, and behavioural processes in mammals ^1–4^. In addition to the primary mammalian clock in the hypothalamic suprachiasmatic nucleus (SCN) ^5^, which is largely regulated by the light/dark period, clock gene transcriptional rhythms are observed in other regions of the brain, including the hippocampus ^6–8^, which remains synchronized to the SCN via a combination of neural and humoral signals ^4,5,9^.

The endogenous corticosteroid hormone cortisol is under circadian control via SCN regulation of hypothalamic-pituitary-adrenal (HPA) axis function. Disruption to this, for example in Cushing’s disease and during steroid therapy -- which elevates cortisol or steroid levels throughout the day, respectively -- is associated with depression, anxiety, and psychosis ^10–12^. Yet, with recent studies showing the importance of the pattern (rather than the level) of steroid availability for improved cognitive and emotional responses ^13^, posits that the rhythmic timing of corticosteroid secretion and therefore activity, is essential to neuropsychiatric health.

Period 1 (*Per1*), a light-entrainable regulator of rhythmic function ^14^, constitutes a key component of the circadian clock machinery targeted by the glucocorticoid receptor (GR: *Nr3c1*) via a hypersensitive glucocorticoid response element (GRE) ^15^. While *Per1* is highly enriched in the SCN and displays robust circadian variation in expression (suppl. Fig. 1), it is resistant to transcriptional disruption by corticosteroids because the SCN is devoid of GR expression ^16^. Similarly, SCN-dependent physiological rhythms, such as locomotor activity and core body temperature are primarily under the control of the SCN transcriptional clock oscillations and are synchronized to the 12-hour light/12-hour dark cycle (suppl. Fig. 2).

GRs are highly enriched in the hippocampus ^16,17^, where their activity displays strong circadian variation in activity, in line with the circadian rise and fall in adrenal corticosteroid secretion, demonstrated by significantly increased GR activity on the *Per1* distal GRE and subsequent *Per1* transcription during the active (Zeitgeber: ZT 14) but not inactive phase (ZT 2) (suppl. Fig. 1; suppl. Fig 3). RNA sequencing (RNAseq) data from rat hippocampi every 4-hours (from ZT 2 – 22) identified 485 differentially expressed genes (DEGs) across time (Fig. 1a; Suppl. Table 1). Of these DEGs, 56 genes (10%) were directly regulated by corticosteroids in an independent experiment where adrenalectomized (ADX) rats were infused with an acute dose of the endogenous corticosteroid corticosterone at ZT 2 (suppl. Fig. 4, Suppl. Table 2).

**Figure 1:**
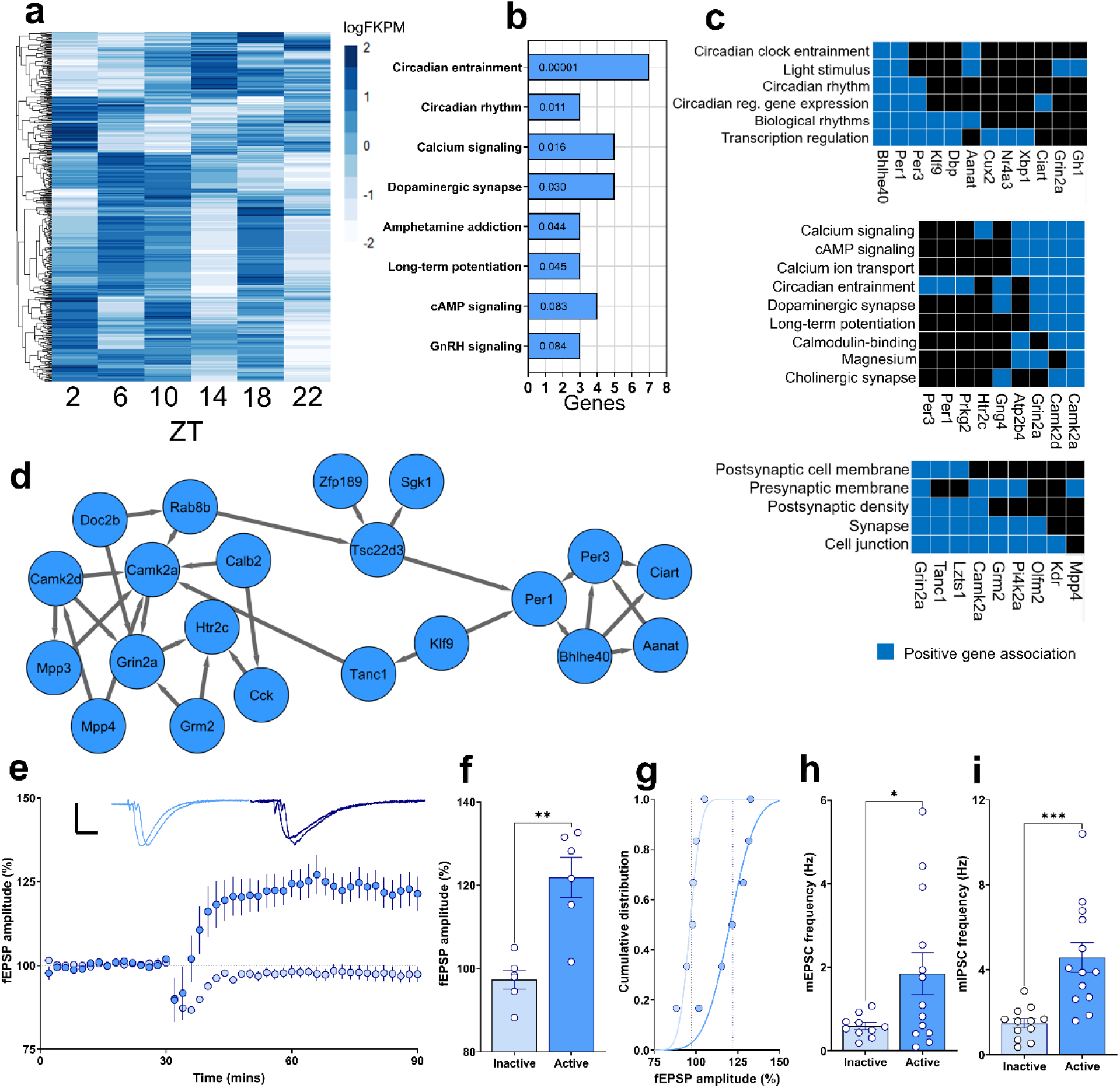
Endogenous regulation of hippocampal function. (**a**) Differentially expressed genes (DEGs; *FDR* < 0.05) across time (*n* = 24, 4/time point). Each row corresponds to a single gene (485 DEGs), each column represents time point. (**b**) Top associated (*P* < 0.085) gene ontology terms among DEGs at ZT 10 (**c**) Functional clustering of DEGs and target pathways at ZT 10. (**d**) Representation of a regulated signaling network enrichment analysis including interacting clock, corticosteroid, and synaptic function DEGs. (**e-g**) Evoked long-term potentiation responses from hippocampal CA1 region neurons recorded for 90 minutes during the early active phase, but not during the early inactive phase (*n* = 5 inactive; *n* = 6 active, *P* = 0.001). (**h-i**) mEPSC and mIPSC measured during the early inactive and early active phase (mEPSC *n* = 10 inactive; *n* = 13 active, *P* = 0.0431: mIPSC *n* = 12 inactive; *n* = 13 active, *P* = 0.0004). Data are mean ± s.e.m. \**P* < 0.05, by unpaired t-test (Fig. 1f, h, i).

In non-stress conditions, endogenous corticosteroid secretion reaches maximal levels near ZT 10, to modulate the awakening response just prior to the start of the active phase ^18^, when many other biological processes are initiated ^19^. During this early period of the active phase, learning and memory processes are more effective, with performance declining as the day progresses ^20–24^. Therefore, we used gene ontology (GO) analysis to identify the biological processes and molecular functions that were enriched in the DEG analysis at ZT 10, relative to the circadian nadir ZT 2 (Fig. 1b). At ZT 10, DEGs included *Per1, Per2, Per3, Bhlhe40, CamkII, Grin2a* and *Grin2c*, known mediators of circadian rhythms and entrainment, as well as modulators of hippocampal synaptic function ^25^. *Per1, Grin2a* and *Grin2b* are directly regulated by acute corticosteroid treatment in ADX rats (Supp Fig. 4, Suppl. Table 2), and GR regulatory sites were identified within or proximal to *Per1, Per2, Bhlhe40, CamkII, Grin2a* and *Grin2b* ^26^. To prepare for the active period, many biological processes have a priming phase, suggesting that genes controlling synaptic function may be regulated by local clock mechanisms in line with the endogenous corticosteroid peak. We clustered the DEGs at ZT 10, by known association (Fig. 1c), discovering functional interactions between core clock gene products – *Per1, Per3, Dbp, Bhlhe40* and *Aanat* - and the known synaptic function genes *Grin2* and *CaMKII* suggesting they are poised to mediate clock-coupled behavioral rhythms to influence learning and memory processes ^27,28^.

Endogenous corticosteroids rapidly regulate GluN2B-NMDAR membrane trafficking through non-genomic mineralocorticoid (MR) signaling ^29^. However, corticosteroids bind to GRs in a circadian pattern, whereas MRs are near maximally occupied at both the circadian nadir and peak due to their higher corticosteroid affinity ^30^. Using a network connectivity map, we identified interactions between enriched genes of corticosteroid -mediated expression, clock gene expression and synaptic function at ZT 10 (Fig. 1d). Genes under circadian and corticosteroid control from independent RNAseq analyses (Fig. 1; Suppl. Fig. 4: *Per1, Klf9, Klf15, Sgk1, Zfp189* and *Tsc22d3)* supports corticosteroid-mediated regulation of clock gene expression ^31,32^. Importantly, peak corticosteroid secretion occurs in preparation for increased neurocognitive activity, which in concert with peak *Per1* clock gene expression in the hippocampus, facilitates action on synaptic plasticity target genes *CamkII* and *Grin2* (Fig. 1d).

Corticosteroid action on memory performance is well-established ^20,33^. However, these studies focusing on the role of corticosteroids have concentrated largely on the response to acute changes in corticosteroids in models of stress ^33,34^. Furthermore, although time-of-day is known to influence memory function ^20,21^, different models and tests have identified conflicting outcomes ^22–24,35–37^. For example, mice and rats learn maze navigation more effectively during the active phase, yet tone-cued fear conditioning in mice is more readily formed during the inactive phase ^35^. Moreover, evidence that tasks with a high cognitive demand can themselves serve as zeitgebers ^38^ suggests that the interaction with time of day is highly stimulus specific. Here we have identified a molecular basis for the improved performance often observed at the start of the active phase. Given that the localization between CaMKII and GluN2B-containing NMDARs at the synapse is critical for long-term potentiation (LTP), a cellular correlate of long-term memory, we hypothesized that this process is under circadian control driven by GR-mediated transcriptional rhythms. In ex vivo hippocampal slices prepared during the early active and inactive periods, LTP was induced only during the early active period (Fig. 1e-i) using a high frequency stimulation protocol; a method used previously to show LTP is modulated by the addition of corticosterone ^39^. The circadian regulation of hippocampal synaptic activity was also reflected with increased frequency, but not amplitude, of spontaneous synaptic events (miniature excitatory postsynaptic currents; mEPSCs and miniature inhibitory postsynaptic currents; mIPSCs) recorded from CA1 neurons (Fig. 1e-i; Suppl. Fig. 5).

Indeed, corticosteroid-mediated memory improvement has been demonstrated in rats, where an acute corticosterone injection during the circadian nadir enhanced memory consolidation in a hippocampal-dependent task ^40^. Chronic corticosteroid exposure, on the other hand, can impair memory in both aversive and non-aversive tasks, and at different times of day ^41,42^. Yet, whether elevated corticosteroid levels, as often seen in major depression, or steroid therapy, mediates memory impairments is difficult to establish in humans, and the mechanisms underlying this relationship remain poorly understood.

Here, we used methylprednisolone (MPL), a long-acting, frequently prescribed synthetic corticosteroid, which is associated with circadian disturbances, memory impairment, and psychiatric disturbances in humans ^43–46^. The binding dynamics of MPL on GRs is distinct from that of endogenous corticosteroid binding, with MPL binding to GRs for > 6-hours ^47^. In agreement with the GC-resistant nature of the SCN, we found that MPL did not alter the circadian expression of core clock genes *Per1* and *Per2*, nor *Avp* (suppl. Fig. 1). Nor did it alter the circadian rhythm of locomotor activity or core body temperature (suppl. Fig. 2), as rhythms remained entrained to the 12-hour light/12-hour dark cycle. However, GRs are highly enriched in the hippocampus, where they are primed to be influenced by endogenous and synthetic corticosteroids. MPL treatment suppressed endogenous circulating CORT (suppl. Fig. 3), but ablated circadian variation in hippocampal GR activity, and instead increased GR activity throughout the 24 hours (including during the inactive period), resulting in increased *Per1* transcription (Suppl. Fig. 3).

Chronic treatment with MPL distinctly dysregulated GR activity and *Per1* transcription. To identify if this altered the hippocampal transcriptome across time, we sequenced the RNA from rat hippocampi every 4-hours (ZT 2 – 22), following five days of MPL treatment. RNAseq analysis identified 512 DEGs across time (Fig. 2a; Suppl. Table 3). Circadian disturbances and memory impairments are common among individuals with major depression or that are receiving steroid therapy. Here, we discovered genes important for memory processing in the hippocampus were dysregulated with MPL treatment.

**Figure 2:**
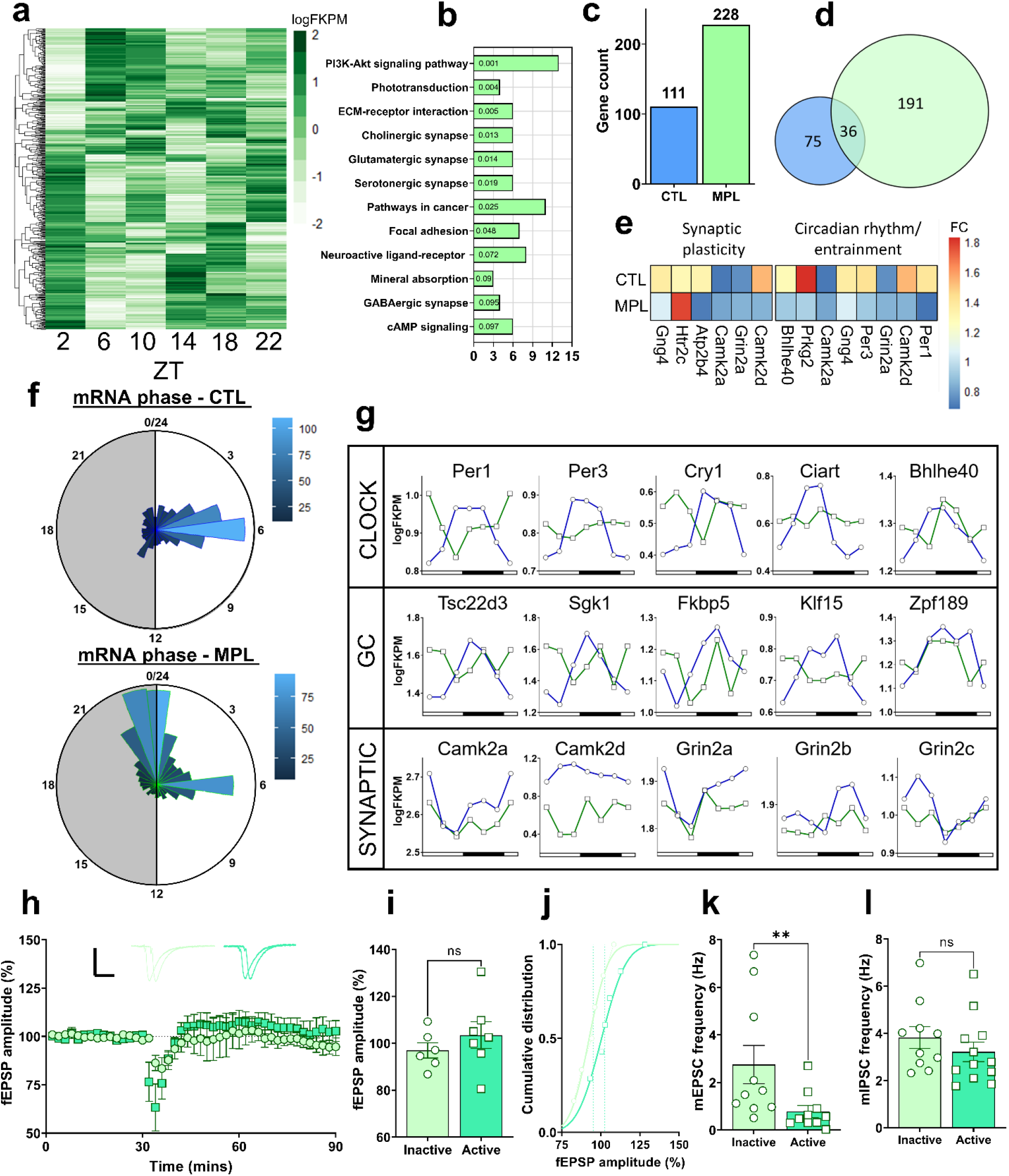
Methylprednisolone disrupts endogenous regulation of hippocampal function. (**a**) Differentially expressed genes (DEGs; *FDR* < 0.05) across time (n=24, 4/time point). Each row corresponds to a single gene (512 DEGs), each column represents time point. (**b**) Top associated (P < 0.1) gene ontology terms of DEGs at ZT 10. (**c**) Total number of DEGs at ZT 10 in CTL and MPL treated groups. (**d**) Overlap of the DEGs at ZT 10 in CTL and MPL. (**e**) Expression of circadian regulating and synaptic function genes at ZT 10 (**f**) Distribution of peak expression of DEGs in CTL and MPL groups. (**g**) RNAseq expression pattern data of clock-regulating, glucocorticoid-target, and synaptic plasticity genes. (**h-j**) MPL treatment prevented LTP responses from hippocampal CA1 region neurons during the early inactive and active phase (n = 5 inactive; n = 6 active, *P* = 0.3692), (**k**) mEPSCs recorded during the early inactive and early active (n = 10 inactive; n = 10 active, *P* = 0.0068) and (**l**) mIPSCs recorded during the early inactive and early active (n = 10 inactive; n = 12 active, *P* = 0.3386). Data are mean ± s.e.m. \**P* < 0.05, by unpaired t-test (Fig. 2i, k), or Mann-Whitney U-test (Fig. 2j).

Following this, we sought to identify if the same functional pathways seen in CTLs were influenced by long-acting corticosteroid treatment. Surprisingly, GO analysis at ZT 10 (Fig. 2b) predicted distinct target pathways compared to equivalent CTL data (Fig. 1b), even though following MPL treatment, additional genes were differentially expressed (Fig. 2c, d; Suppl. Table 4). Interestingly, no circadian entraining pathways were identified with GO analysis, suggesting a loss of hippocampal rhythm, and although synaptic plasticity pathways were identified at ZT 10 with MPL treatment, the DEGs identified at these times were not evident in CTLs (Suppl. Table 5). With differential pathway activity between CTL and MPL groups, we quantified the expression of circadian rhythm and entrainment genes, as well as synaptic plasticity target genes at ZT 10 in CTL and MPL groups (Fig. 2e). Following the detection of a dissociation between clock regulating and synaptic plasticity genes at ZT 10, we sought to determine the peak amplitude of all DEGs across the day (Fig. 2f; Suppl. Table 6) ^48^. We found that most DEGs in CTL exhibited peak expression levels between ZT 5-7 (Fig. 2f), whereas following steroid treatment, most DEGs peaked between ZT 22-1 (Fig. 2f). Subsequent to the shift in peak gene expression, we characterized gene expression across time of known clock-regulating, corticosteroid-target, and synaptic plasticity-associated genes (Fig. 2g). The striking pattern differences, and peak expression profiles of genes regulating clock and synaptic plasticity processes following MPL treatment led us to investigate a functional output of this disruption. In CTLs, LTP was robust during the early active period, as was the frequency of mPSCs (Fig. 1e-i). Using the same HFS protocol, but this time in rats that received five days of MPL treatment, LTP could not be induced in either the early active or inactive period (Fig. 2h-i). Remarkably, MPL treatment increased the frequency, but not amplitude, of mEPSCs recorded during the early inactive period, an inversion of the pattern observed in controls. This was not evident in the frequency of mIPSC events, instead eliminating circadian variation, suggesting MPL disrupts the excitatory/inhibitory (E-I) mPSC balance (Fig. 2j-k; Suppl. Fig. 5). Interestingly, alterations to the E-I balance in the forebrain has been implicated in several psychiatric disorders including Alzheimer’s disease ^49,50^. Additionally, early-life stress, accelerates developmental shifts in the E-I balance ^51^, emphasizing the role of corticosteroid signaling in mediating the regulation of this phenomenon.

CaMKII is essential for modulating long-term potentiation and spatial memory formation ^52^, and following the striking disruption to the dynamic expression of both NMDA receptor and CaMKII subunits, we show that hippocampal long-term potentiation and mPSCs were disrupted following MPL treatment. Again, using the same HFS protocol that was sensitive to regulation by time-of-day (Fig. 1e-i), MPL treatment abolished the formation of LTP during the early active period, when it could be readily elicited in CTLs (Fig. 3a-c). Furthermore, in CTLs, LTP induction could be similarly blocked by application of the GluN2B selective NDMAR antagonist, Ro-25-6981 (Fig. 3d-f). This suggests that ablation of the circadian regulation of NMDARs and CaMKII results in disruption of downstream plasticity processes by prevention of their association at a time critical to non-aversive learning. Consistent with our data demonstrating corticosteroid sensitivity, NMDARs are critical in mediating the dendritic atrophy and synaptic dysfunction caused by chronic stress ^53,54^ with GluN2B playing a pivotal role ^55^.

**Figure 3:**
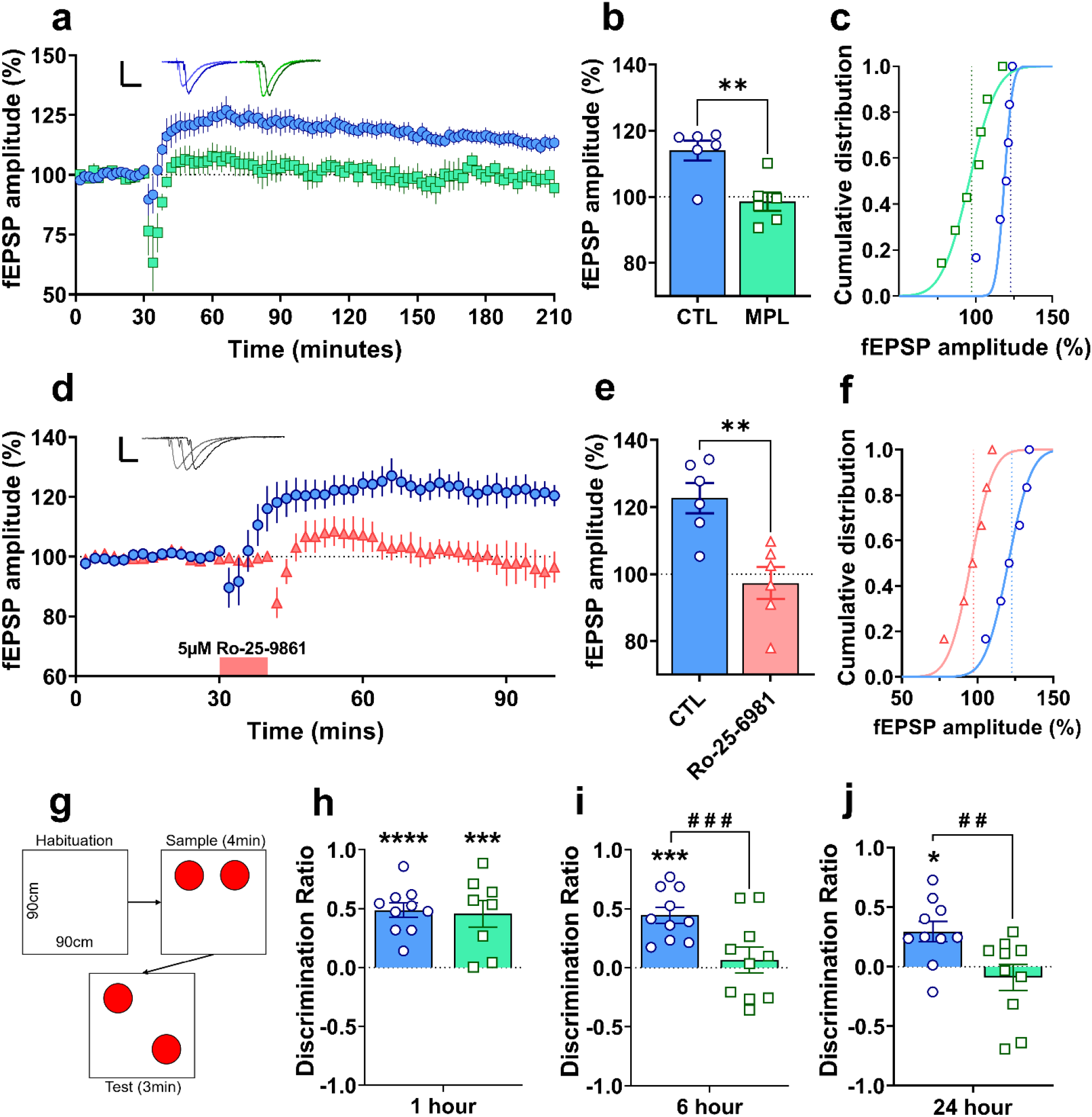
Methylprednisolone inhibits NMDAR-dependent LTP and impairs intermediate and long-term hippocampal-dependent memory formation. (**a-c**) LTP recorded during the early active phase for 3 hours post-high frequency stimulation in the hippocampal slice from CTL and MPL treated rats and quantified during a 30-minute epoch 2.5 hours post-HFS (n = 6 CTL active; n = 6 MPL active, *P* = 0.0037). (**d-f**) LTP recorded following 5 µM Ro-25-6981 application and HFS stimulation (n = 6 CTL active; n = 6 Ro-25-6981 active, *P* = 0.0031). (**g**) Schematic of Object Location Memory (OLM) task. (**h**) OLM 1hr post acquisition phase in CTL and MPL treated rats. *F*_1,32_ = 59.52, *P* < 0.0001, for learning, by two-way ANOVA. \*\*\*\**P* < 0.0001. (**i**) OLM 6hr post acquisition phase in CTL and MPL treated rats. *F*_1,36_ = 15.87, *P* = 0.0003, for memory, and *F*_1,36_ = 8.727, *P* = 0.0055, for treatment, by two-way ANOVA. \*\*\**P* = 0.0001, ^###^*P* = 0.0010, Tukey’s multiple comparisons. (**j**) OLM 24hr post acquisition phase in CTL and MPL treated rats. *F*_1,36_ = 7.646, *P* = 0.0089, for treatment, by two-way ANOVA. \**P* = 0.0245, ^##^*P* = 0.0021, Tukey’s multiple comparisons. Data are mean ± s.e.m. \**P* < 0.05.

Following the robust effects of steroid treatment on the hippocampal transcriptome and synaptic physiology, we investigated whether MPL-induced disruption of NMDA receptor function had a consequent impact on hippocampal-dependent memory formation. After five days of MPL treatment, using a hippocampal-dependent memory task, where training and testing were carried out during the early active period, short-term memory retrieval (1-hour post-acquisition phase) was unaffected (Fig. 3h). However, memory retrieval was impaired at intermediate (6-h) and long-term retrieval (24-h), post-acquisition phase (Fig. 3i+j) which was not driven by changes in exploration time (Suppl. Fig. 6). These data suggest that MPL treatment inhibits long-term, GluN2B-NMDAR dependent memory, but not short-term memory formation ^56,57^.

## Discussion

Corticosteroids are widely used in clinical medicine to relieve signs and symptoms of many inflammatory and autoimmune disorders ^58,59^. However, in addition to this, their use is often reported with cognitive and psychiatric symptoms – inducing a range of psychiatric adverse effects including depression, mania, and cognitive and memory impairment. This relationship has been known for decades, yet the mechanisms of corticosteroid-induced psychiatric disturbances and their clinical management has been poorly described.

Circadian rhythms are routinely disrupted across several psychiatric disorders ^60–62^. Adrenal corticosteroid secretion, governed largely by the circadian control of the master clock (SCN) on the HPA axis, is often hyperactive in depressed patients. Yet, identifying the causal relationship between dysregulated HPA axis function (i.e., elevated steroid availability throughout the day) and neuropsychiatric disorders is difficult to describe in humans.

In this study, and using rats to limit external influences, we demonstrate that five days of corticosteroid treatment disrupts the rhythmic activity of hippocampal function through GRs, which in turn mediates NMDAR-dependent synaptic dysfunction and subsequent impaired memory. This model is supported by several lines of evidence: (1) Corticosteroids do not influence the master clock (SCN), or light/dark entraining behaviours due to a lack of GRs present; (2) The hippocampus is rich in GRs, and in the presence of corticosteroids, binds to the GRE on the essential clock gene *Per1* to mediate hippocampal activity in competition with central clock-mediated control; (3) Synthetic corticosteroid treatment eliminates the circadian binding of corticosteroid on GRs in the hippocampus to influence clock gene expression and destabilize clock entrained NMDAR/CamkII complexes; and (4) Synthetic corticosteroid treatment blocks LTP, a cellular correlate of long-term memory, by disrupting NMDAR mechanisms processes to impair long-term, but not short-term, hippocampal-dependent memory.

There are two regulators in normal chronophysiology. On the one hand, the SCN is the master regulator of circadian timing, entraining behaviours such as feeding, sleep, and memory processes to ∼24hr ^63^. On the other hand, extra-SCN oscillators, such as those in the hippocampus and other forebrain regions are entrained by SCN-dependent neural and humoral inputs ^64^. Pertinent to this study, one of the more powerful humoral signals is from adrenal corticosteroids ^65^, which exhibit a robust circadian secretory rhythm that peaks upon wakening ^2^ in preparation for the neurocognitive activities that ensue. Synthetic long-acting corticosteroids, such as MPL, are therefore poised to mediate influence over these endogenous systems in the presence of the master clock.

Indeed, corticosteroids exert vast and variable effects on the hippocampus including memory processing ^66^. Here, we found that the SCN remained protected from MPL-mediated disruption, similarly to Dexamethasone ^67^. However, extra-SCN oscillators, such as in the GR-rich hippocampus, are sensitive to corticosteroid interference. For example, the identification of the GRE on the *Per1* gene and corticosteroid-induced transcription, suggests that extra-SCN oscillators can be disturbed by corticosteroids, and may account for the loss of circadian rhythms often seen in hypercortisolaemic states of Cushing’s disease and syndrome ^68^.

It is well established that corticosteroids are secreted from the adrenal gland in a circadian pattern with underlying ultradian pulses, and common to several neuropsychiatric disorders are HPA axis hyperactivity or steroid therapy, and cognitive and memory impairments ^46,69^. We found that five days of corticosteroid treatment (which elevated the level of circulating corticosteroids and subsequently removed the circadian and ultradian pattern of secretion) disrupted the circadian expression of NMDAR and CAMKII subunits, both of which are essential mediators of neural plasticity ^70^, a process that is causally linked with the successful encoding of mnemonic information ^71^. Further to this, CAMKII is essential for coupling time-of-day to behavioural rhythms ^27^ and improved memory performance ^72^.

Consistent with our findings, other synthetic corticosteroids, particularly Dexamethasone, can impair memory ^73^. However, translational studies often assess memory performance during the early inactive period, when endogenous circulating corticosteroids are low, therefore not time-relevant to the biological rhythms ^74^ or peak cognitive performance of the participants. Similarly, many rodent studies assess both hippocampal synaptic plasticity processes and memory performance during the animals’ inactive period, leading to the somewhat incomplete conclusion that effective memory consolidation requires a strong stress-association. These tests use aversive or fear conditioning-based tasks such as inescapable foot shock, which induces a robust stress response in the animal, resulting in similar elevated glucocorticoid levels to those circulating at the onset of the active phase. Consistent with our interpretation that elevated glucocorticoids and the consequent effects on the molecular physiology of brain, acute administration of corticosteroids has been demonstrated to enhance memory consolidation during a non-aversive memory task performed during rats’ inactive phase. ^75,76^. The specificity of corticosteroid, duration of action, as well as the timing and type of experimental testing, may contribute to differences in corticosteroid-mediated memory formation in the literature. Importantly in this study, we have identified a mechanism by which neurocognitive decline is mediated by a commonly prescribed synthetic corticosteroid. However, how various corticosteroids (endogenous and synthetic), contribute to such differential outcomes on hippocampal function warrants further investigation.

In line with this, our ex vivo physiology data suggest that time of day is a potent modulator of neuronal activity and plasticity processes in CA1 neurons. Crucially, we show that this circadian regulation is modified by long-acting corticosteroid treatment, disrupting the dynamic control of miniature post-synaptic currents (mPSCs) and preventing LTP induction. These data support the notion that circadian and ultradian corticosteroid fluctuations are essential for maintaining the rhythmic activity of the hippocampus via dynamic transcription of target genes. This has major physiological consequences considering the importance of mPSCs in the maintenance of synaptic connections and dendritic spines ^77^, as long-term alterations to these dynamics could result in critical changes to hippocampal circuitry and function, as demonstrated by the prevention of hippocampal-dependent memory formation.

In summary, a large proportion of patients prescribed corticosteroids report cognitive decline and memory impairment ^43^. Our data reveal a GR-mediated pathway that underlies the circadian regulation of hippocampal-dependent memory formation which is vulnerable to long-acting corticosteroid treatment. Currently, there are no clinical guidelines for treating corticosteroid-induced psychiatric adverse effects. It is, perhaps, quite surprising that the brain-specific effects of such treatments have had such little scientific investigation given their broad application, widespread clinical use and adverse psychiatric effects ^45,78,79^. Advancing our knowledge of corticosteroid-mediated regulation of hippocampal function will take us a step closer to understanding the mechanisms underpinning several prevalent major mental illnesses and suggest novel methods for prevention and intervention of corticosteroid (endogenous and synthetic)-induced disorders.

## Abbreviations

AVP: Vasopressin
BHLHE40: Basic helix-loop-helix e40
CaMKII: Calcium/calmodulin-dependent protein kinase type II
CORT: Corticosterone
DBP: D-box protein
DEGs: Differentially expressed genes
GluN2B: Glutamate [NMDA] receptor subunit 2B
GO: Gene ontology
GR: Glucocorticoid receptor
GRE: Glucocorticoid response element
GRIN2: Glutamate ionotropic receptor NMDA type subunit 2
HFS: High frequency stimulation
KLF15: Kruppel like factor 15
LTP: Long term potentiation
mEPSC: miniature Excitatory Post-synaptic Currents
mIPSC: miniature Inhibitory Post-synaptic Currents
MPL: Methylprednisolone
MR: Mineralocorticoid receptor
NMDAR: N-methyl-D-aspartate receptor
PER1: Period 1
PER2: Period 2
PER3: Period 3
SCN: Suprachiasmatic nucleus
SGK1: Serum/Glucocorticoid regulated kinase 1
TSC22D3: Glucocorticoid Induced Leucine Zipper
VEH: Vehicle
ZFP189: Zinc finger protein 189
ZT: zeitgeber time

---

**Supplementary Fig. 1:**
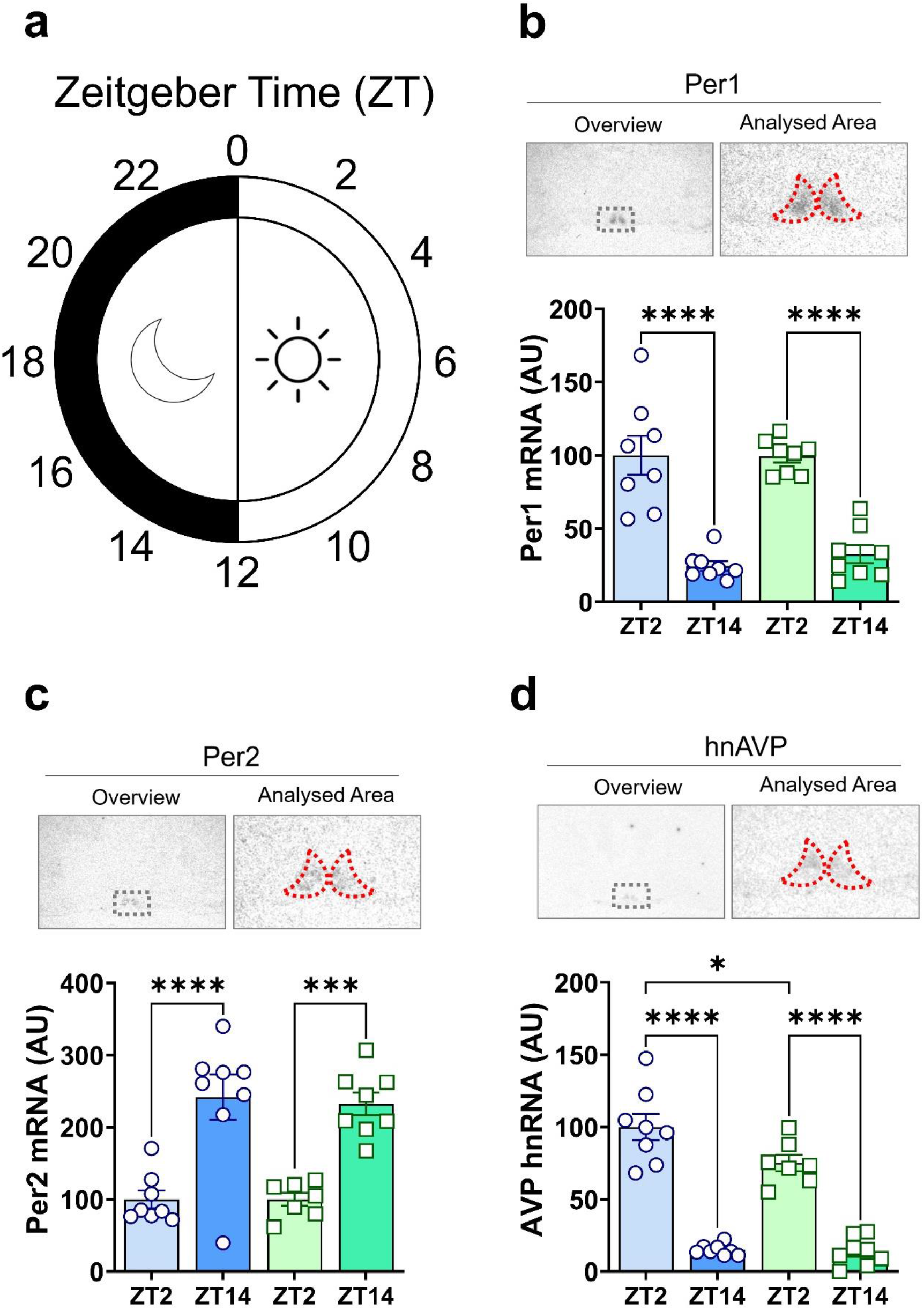
Clock gene expression in the SCN is protected from methylprednisolone treatment. (**a**) Schematic showing experimental timing relative to light cycle. **(b)** Period 1 mRNA expression in the SCN in CTL and MPL treated rats. *F*_1,28_ = 83.97, *P* < 0.0001, for time by two-way ANOVA. \*\*\*\**P* < 0.0001, Sidak’s multiple comparisons. (**c**) Period 2 mRNA expression in the SCN of CTL and MPL treated rats. *F*_1,27_ = 48.41, *P* < 0.0001, for time by two-way ANOVA. \*\*\*\**P* < 0.0001, \*\*\**P* = 0.0001, Sidak’s multiple comparisons test. (**d**) AVP hnRNA expression in the SCN of CTL and MPL treated rats. *F*_1,27_ = 166.1, *P* < 0.0001, for time by two-way ANOVA. *F*_1,27_ = 5.283, *P* = 0.0295, for treatment by two-way ANOVA. *F*_1,27_ = 4.316, *P* = 0.0474, for an interaction. \*\*\*\**P* < 0.0001, \**P* = 0.0307. Sidak’s multiple comparisons test. Data are mean ± s.e.m. \**P* < 0.05.

**Supplementary Fig. 2:**
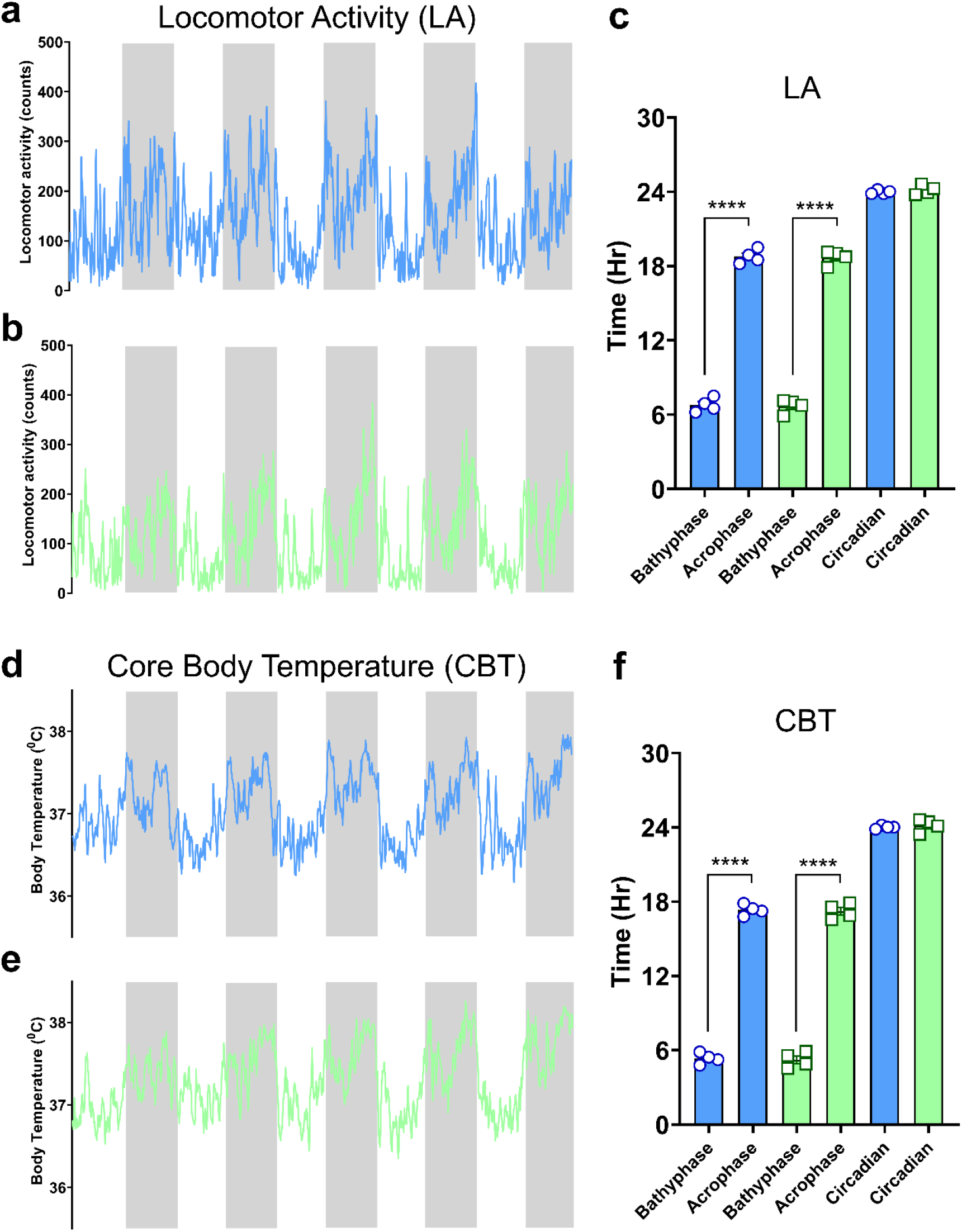
Locomotor activity and core body temperature are unaffected by methylprednisolone treatment. Example locomotor activity in a (**a**) CTL and (**b**) MPL treated rat. (**c**) CTL and MPL treated rats maintained rhythmic locomotor activity (*F*_1,12_ = 1801, *P* < 0.0001: CTL; *P* < 0.0001, MPL; *P* < 0.0001). Example core body temperature in a (**d**) CTL and (**e**) MPL treated rat. Like activity, (**f**) MPL treatment did not influence core body temperature rhythm (*F*_1,12_ = 1964, *P* < 0.0001: CTL; *P* < 0.0001, MPL; *P* < 0.0001). Two-way ANOVA with Sidak’s multiple comparisons test. Data are mean ± s.e.m. \**P* < 0.05.

**Supplementary Fig. 3:**
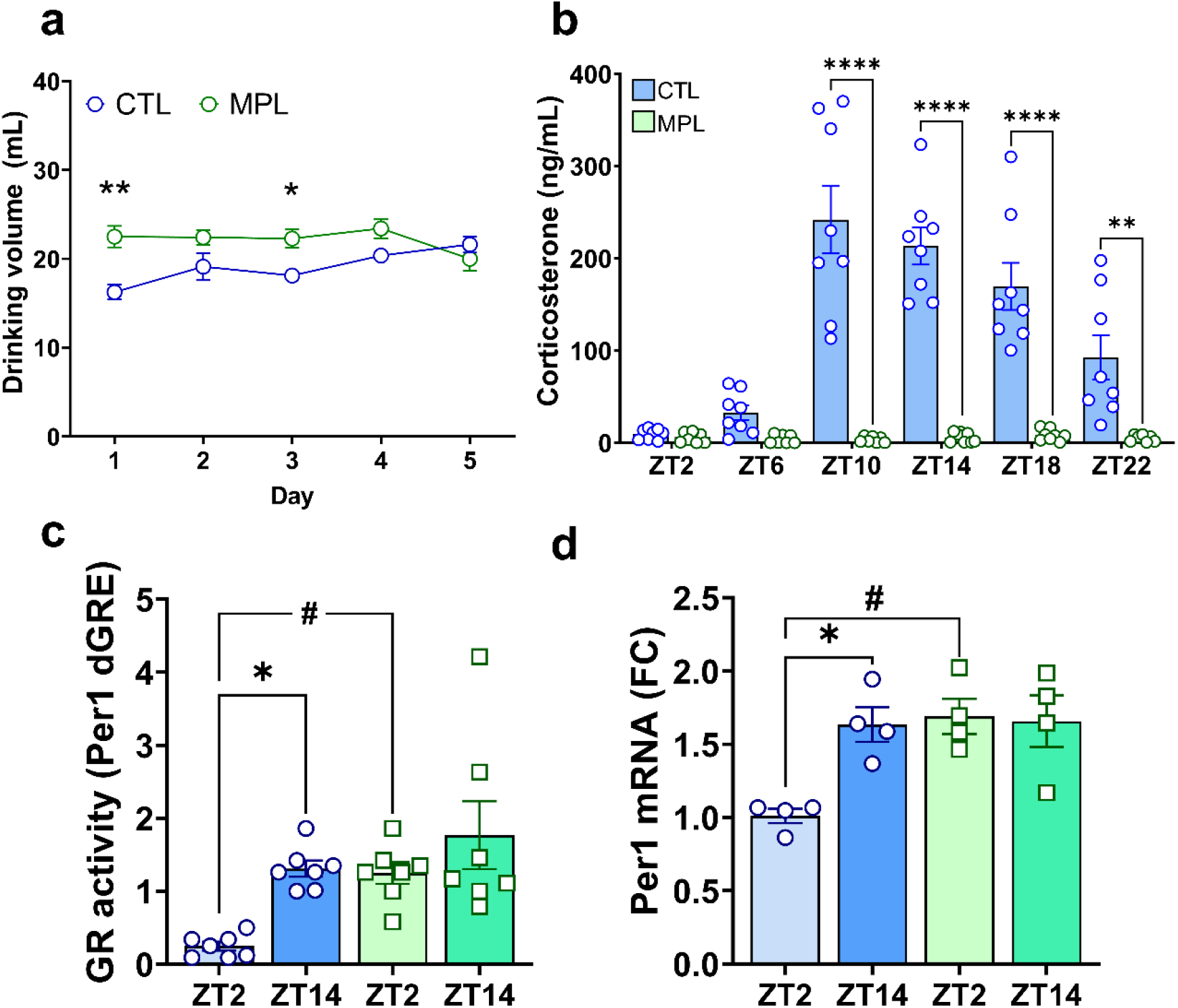
Methylprednisolone suppresses endogenous circulating CORT and alters hippocampal GR binding and mRNA expression. (**a**) Drinking volumes from CTL and MPL treated rats across the treatment length. No effect of day was observed on consumption (*F*_3.203,41.64_ = 1.647, *P* = 0.1905), but there was an effect of treatment (*F*_1,14_ = 18.48, *P* = 0.0007) with an increase in total drinking volume in MPL rats above controls (Day 1: P=0.0052; Day 3: P=0.0339, Sidak’s multiple comparisons). (**b**) Methylprednisolone in drinking water was sufficient to inhibit circulating corticosterone in rats. *F*_5,81_ = 16.98, *P* < 0.0001, for time, and *F*_1,81_ = 160.6, *P* < 0.0001, for treatment by two-way ANOVA. \*\*\*\**P* < 0.0001, ^**^*P* = 0.0026, Sidak’s multiple comparison. (**c**) Hippocampal GR activity on Per1 dGRE. *F*_1,24_ = 9.803, *P* = 0.0045, for time, and *F*_1,24_ = 8.339, *P* = 0.0081, for treatment by two-way ANOVA. \**P* = 0.0314, ^#^*P* = 0.0458, Tukey’s multiple comparison. (**d**) Hippocampal Per1 mRNA expression. *F*_1,12_ = 5.593, *P* = 0.0357, for time, and *F*_1,12_ = 7.925, *P* = 0.0156, for treatment by two-way ANOVA. \**P* = 0.0186, ^#^*P* = 0.0106, Tukey’s multiple comparison. Data are mean ± s.e.m. *P<0.05.

**Supplementary Fig. 4:**
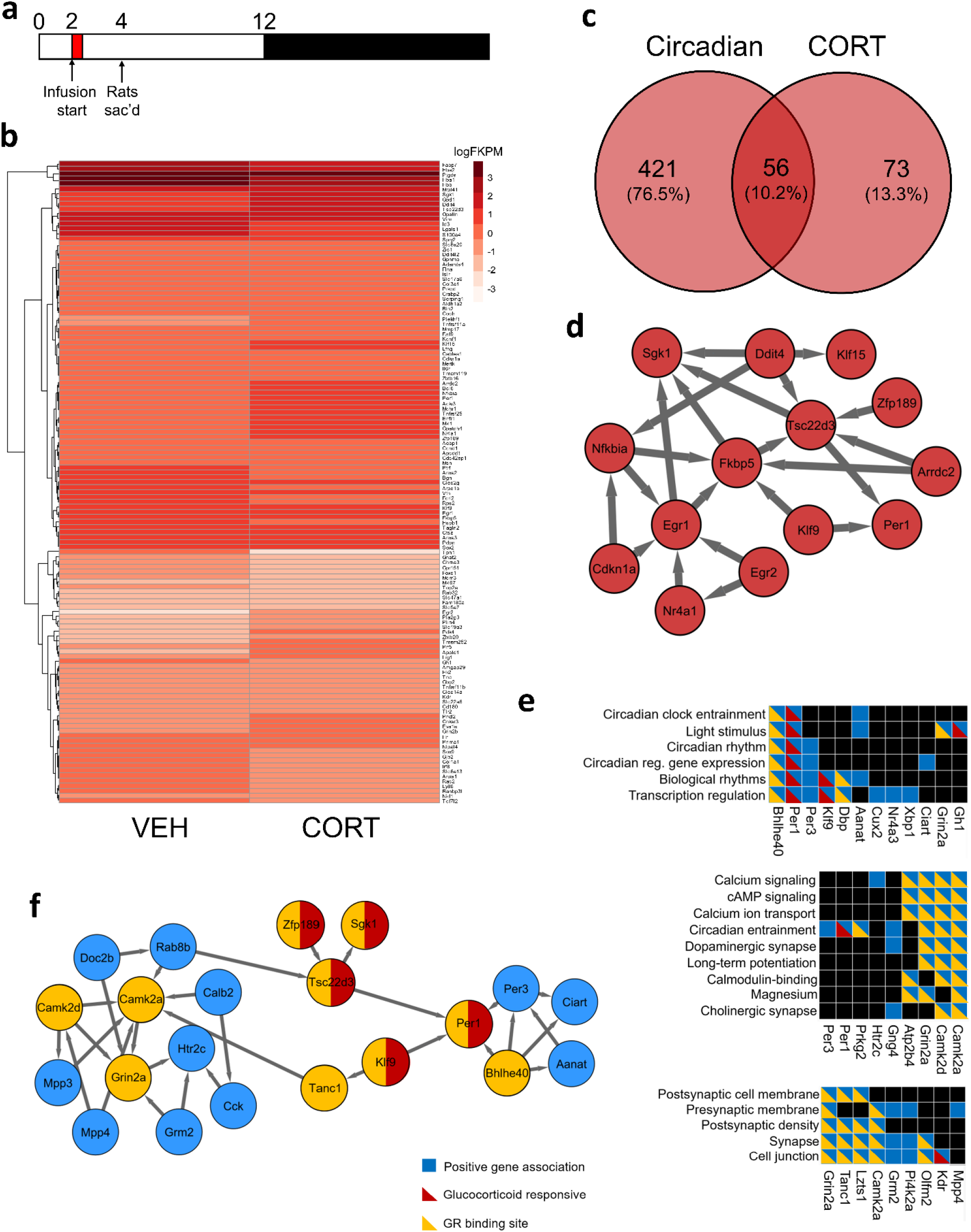
Hippocampal RNAseq of Vehicle and CORT infused ADX rats. (**a**) Schematic of timing of VEH or CORT infusion and hippocampi collection. (**b**) Heatmap of DEGs from vehicle infused and CORT infused ADX rats (n = 4 vehicle, n = 4 CORT). (**c**) Of total DEGs in Fig. 1, 56 (10.2%) DEGs are regulated by glucocorticoids. (**d**) Representation of a regulated signaling network enrichment analysis of glucocorticoid responsive genes that are under circadian control. (**e**) Overlap of functional clustering of DEGs at ZT 10 from Fig. 1, with identified glucocorticoid responsive (red) and associated GR binding site (yellow). (**f**) Representation of a regulated signaling network enrichment analysis genes that are under circadian control (blue), glucocorticoid responsive (red), and/or contain a GR binding site (yellow).

**Supplementary Fig. 5:**
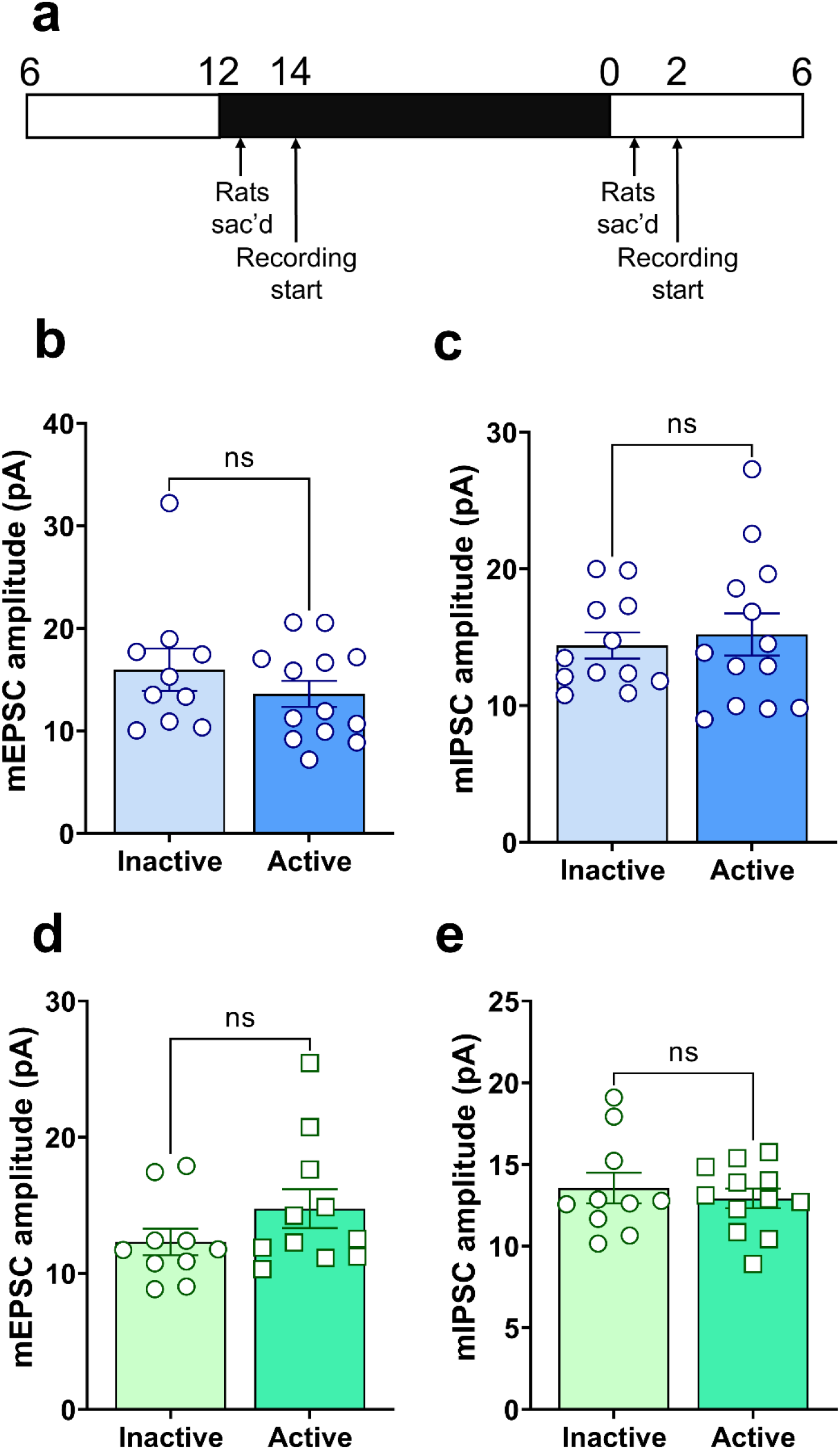
mEPSC or mIPSC amplitudes were not influenced by time of day or methylprednisolone treatment. (**a**) Schematic of timing of electrophysiology experiments. (**b**) Time of day did not affect the amplitude of mEPSCs (n = 10 inactive; n = 12 active, *P* = 0.3157) nor (**c**) mIPSCs (n = 12 inactive; n = 13 active, *P* = 0.6671). MPL treatment did not influence time of day (**d**) mEPSCs (n = 10 inactive; n = 11 active, *P* = 0.1812) or (**e**) mIPSCs (n = 10 inactive; n = 12 active, *P* = 0.5645).

**Supplementary Fig. 6:**
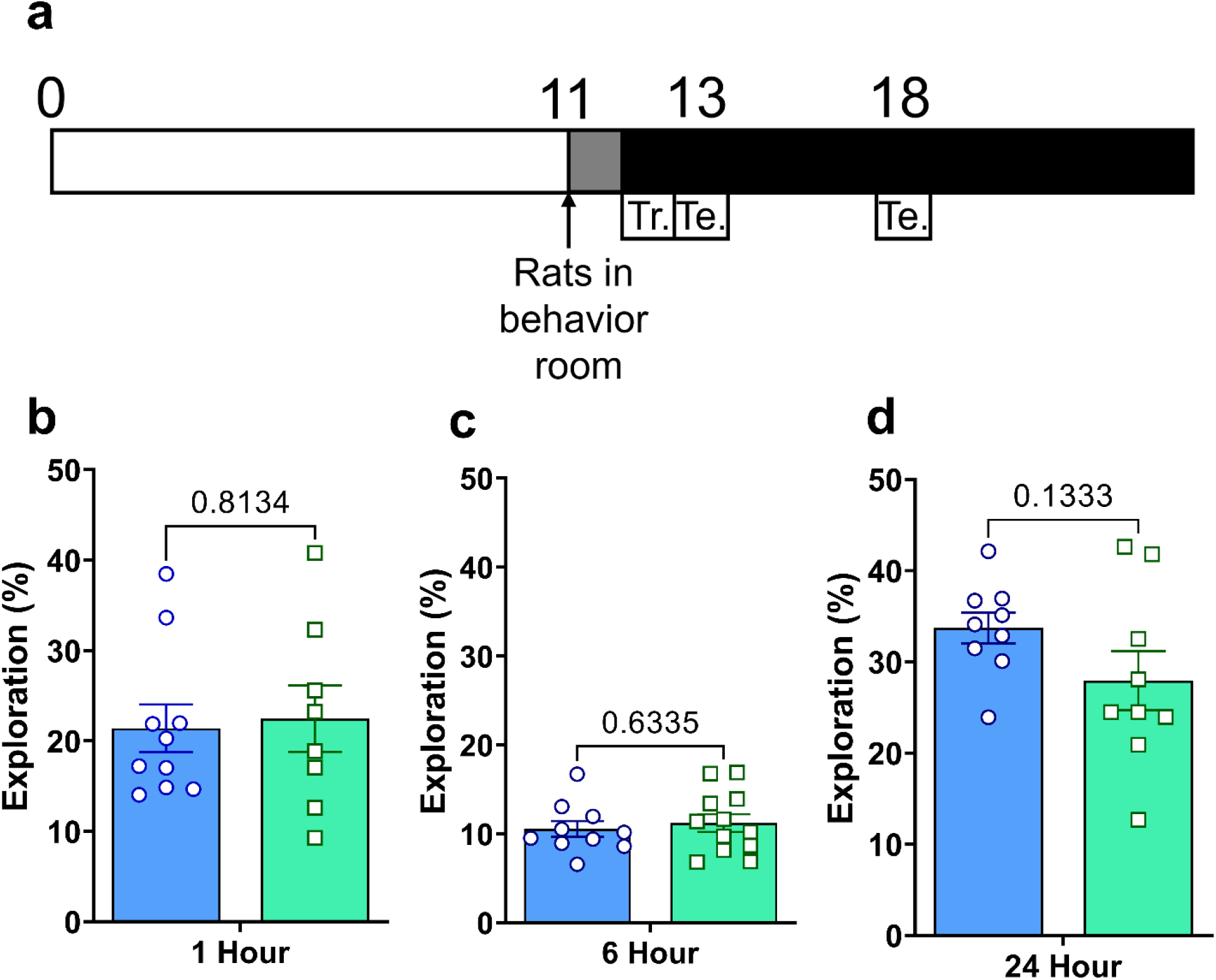
Methylprednisolone did not influence exploration during object location memory testing. (**a**) Schematic of object location experimental timing. (**b**) Exploration of objects in CTL and MPL treated rats 1hr post sample phase (n = 10 CTL; 8 MPL, *P* = 0.8134) (**c**) Object exploration in CTL and MPL treated rats 6hr post sample phase (n = 10 CTL; 12 MPL, *P* = 0.6335). (**d**) Object exploration in CTL and MPL treated rats 24hr post sample phase (n = 9 CTL; 9 MPL, *P* = 0.1333), Data are mean ± s.e.m. Tr; Training. Te; Test.

## Methods

Adult male Lister hooded rats (250-350g, 9-11 weeks) (Envigo, UK) were used in all experimental procedures. Animals were maintained in standard housing conditions under a 12:12 gradual light/dark cycle in sound-attenuating rooms. A red light was used during the night phase to allow researchers to manipulate the animals. During the night phase, no external light could enter the room when opening the door. Food and water were available *ad libitum*. Throughout all experimental procedures, the same researcher took care of the animals to limit stress-induced effects and was blinded to treatment groups. All procedures were carried out in accordance with the UK Animals (Scientific Procedures) Act 1986 under PPL 30/3114 and PIL I04092F5F.

### Adrenalectomy and jugular vein cannulation

For studies identifying glucocorticoid-specific control of hippocampal gene transcription (Suppl. Fig. 4), rats received balanced anesthesia using veterinary isoflurane (Merial Animal Health, Woking, UK) prior to bilateral adrenalectomy and cannulation of the right jugular vein for infusion of corticosterone. Rats recovered for 5 days post-surgery on 15 µg/mL corticosterone in 0.9% saline drinking solution to maintain isotonic levels. This solution was replaced 12 hours prior to experiments with 0.9% saline to ensure washout of circulating corticosterone.

### Corticosterone infusion

Rats received either a 30-minute infusion of corticosterone (0.75 mg/mL Corticosterone-2-hydroxypropyl-*β-*cyclodextrin; Sigma-Aldrich, Gillingham, UK) dissolved in sterile 0.45% (w/v) NaCl, or sterile 0.45% (w/v) NaCl at ZT2. A New Era NE-1800 computer-driven infusion pump (World Precision Instruments, Aston, UK) delivered 1 mL/h for 30 minutes via the indwelling jugular cannula. Rats were killed 2 hours post infusion start at ZT4 (Suppl. Fig. 4).

### Methylprednisolone treatment

For all studies assessing the role of Methylprednisolone (MPL), rats (randomly assigned to treatment) were given Methylprednisolone (Solu-Medrol; Pharmacia Ltd, Sandwich, UK) 1 mg/mL ad libitum in drinking water for 5 days (chronic), prior to the start of all further procedures. Ad libitum access to MPL in drinking water was designed to limit any external stimuli that may act as a zeitgeber^80^, and also represents greater than 1% of the human population that are prescribed oral glucocorticoid treatment^45^. At this dose, no difference in fluid consumption was observed across days and circulating endogenous corticosteroids (measured at the end of 5 days treatment) were suppressed (Suppl. Fig. 3).

### Radioimmunoassay

Using an automated gamma counter (PerkinElmer, US), a corticosterone radioimmunoassay measured endogenous levels in blood samples collected immediately following rapid decapitation. An 11-point standard curve of known corticosterone concentrations was prepared in B-buffer (25mM tri-sodium citrate, 50mM sodium dihydrogen orthophosphate, 1mg/mL bovine serum albumin: pH3). Plasma obtained was diluted in triplicate at a ratio of either 1:10 or 1:50 in B-buffer, respectively. A specific corticosterone antibody (kindly provided by G. Makara, Institute of Experimental Medicine, Budapest, Hungary) was diluted at a ratio of 1:50 in B-buffer and 50μl added to 100uL standards, unknown samples, and Quality Control (QC20 and QC100) tubes. Tracer (Izotop, Institute of Isotopes, Hungary) was diluted in B-buffer to give total counts of 3750cpm in 50μl and added to all tubes (50μl/tube). Tubes were incubated overnight at 4°C. Charcoal suspension (5g charcoal added to 0.5g Dextran T70 dissolved in 1L B-buffer) was prepared and 500μl added to all tubes and briefly vortexed. Blocks were centrifuged at 4000rpm at 4°C and the resulting supernatant aspirated off. Unknown samples were determined from interpolation of the standard curve.

### In situ hybridisation histochemistry

Whole brains were cryosectioned into coronal 12 μm sections, and thaw mounted on gelatin/chrome alum-coated slides. The location of the SCN was determined according to coordinates in the rat brain atlas^81^. ^35^S 3’- end-labeled deoxyoligonucleotide complementary to transcripts encoding *Per1 (*5’- CTC TTG TCA GGA GGA ATC CGG GGA GCT TCA TAA CCA GAG TGG ATG -3’), *Per2 (*5’- GTG GCC TTC CGG GAT GGG ATG TTG GCT GGG AAC TCG CAC TTT CTT -3’) and *hnAVP* (5’- GCA CTG TCA GCA GCC CTG AAC GGA CCA CAG TGG TAC -3’) were used. The *in situ* hybridization protocol has been previously described in detail^82^. Briefly, sections were fixed in 4 % (w/v) formaldehyde for 5 minutes and incubated in saline containing 0.25 % (v/v) acetic anhydride and 0.1 M triethanolamine for 10 minutes. Sections were then dehydrated in ethanol, delipidated in chloroform, and partially rehydrated. Hybridization with with a total count of 1 × 10^6^ cpm was performed overnight at 37 **°**C in 45 μL of hybridization buffer under Nescofilm (Bando Kagaku, Osaka, Japan). After hybridization, sections were washed 4 times with SSC (150 mM NaCl and 15 mM sodium citrate) for 1 hour at 65 °C and for an additional hour with two changes of SSC at room temperature. Hybridized sections were exposed for autoradiography (Hyperfilm, Amersham, Bucks, UK) for 1 week. The amount of bound probe was analysed in comparison to ^14^C-labelled standards (Amersham, Bucks, UK) using image analysis software (NIH Image 1.6.2, W.Rasband, NIH, Bethesda, MD, USA). The obtained results were represented in arbitrary units setting the mean optical density (OD) obtained from sham operated rats.

### Chromatin immunoprecipitation (ChIP)

Hippocampi were fixed for ChIP and processed to chromatin in sodium dodecyl sulfate (SDS) lysis buffer [2% SDS, 10 mM EDTA, 50 mM Tris-HCl (pH 8.1)] as previously described^83^. Chromatin was sheared with a Sonifier 450 (Branson Ultrasonics, Danbury, CT) using 4× 10-second pulses at 10% output, and then cleared of cellular debris by centrifugation. For each IP, chromatin was diluted 1:10 in ChIP dilution buffer [167mM NaCl, 16.7mM Tris-HCl (pH 8.1), 1.1% Triton X-100, 1.2mM EDTA, 0.01% SDS] supplemented with cOmplete protease inhibitor (Sigma). Reactions were immunoprecipitated overnight at 4°C with 2µg anti-GR M-20X (Santa Cruz Biotechnology, US) or rabbit non-immune serum (2µg of sc-2027; Santa Cruz, USA) for the negative control. GR–DNA complexes were collected onto protein A–conjugated Dynabeads (Invitrogen, Paisley, UK) and washed to remove nonspecific binding. Purified DNA was resuspended in nuclease free water (Ambion, Huntington, UK).

PCR primers (forward: 5′- CCAAGGCTGAGTGCATGTC -3′; reverse: 5′- GCGGCCAGCGCACTA -3′) were designed to amplify across a previously described GRE^15,83^ in the rat Period 1 gene promoter. Samples were amplified with Sybr Green master mix (Applied Biosystems) in accordance with the manufacturer’s instructions. GR binding for each sample was calculated relative to 1% input chromatin taken from each individual sample, using the %Input method, described in (ThermoFisher Scientific, http://bit.ly/ChIPAnalysisTFS).

### RNA and qPCR

Animals were anesthetized with isoflurane in the animal facility, and eight were killed every 4 hours (six time points). The brain was quickly extracted, and the hippocampus was removed and rapidly frozen in liquid nitrogen. The time between decapitation and sample freezing was < 1 min to limit RNA degradation. The collection of the hippocampal tissue per time point was < 20 min (10 min either side of the hour mark).

Total RNA was extracted from frozen whole hippocampi using the miRNeasy total RNA extraction kit protocol (Qiagen, US) following the manufacturer’s guide and included a DNase digestion step. Samples were stored at -80°C. All samples were assessed for RNA quality and quantity using a Nanodrop (ThermoScientific, US). Samples sent for RNA sequencing were further assessed for RNA integrity on the 2200 Tapestation system (Agilent, US). Sequenced samples had >8.0 RIN score.

Each PCR reaction contained 1µL of cDNA, total volume 10µL. qPCR runs consisted of an initial 95°C holding stage for 20 seconds, followed by 40 cycles of 95°C (1 second) and 60°C (20 second), followed by a melt curve step, consisting of 40 cycles of 95°C (15 seconds) and 60°C (1 minute), with a final denaturing step of 95°C (15 seconds) using a StepOnePlus PCR machine (Applied Biosystems, Life Technologies, UK).

### Whole genome RNA-sequencing and analyses

RNA sequencing of hippocampal tissue was carried out using TruSeq Stranded Total RNA kit and protocols (Illumina, US). A 1ug aliquot of total RNA from each hippocampus was prepared for RNA sequencing. 48 samples total. First, bioanalyzer traces were carried out on all samples for RNA quantity and quality check. RIN scores above 8.0 were deemed high enough quality to take through to RNA sequencing. Following library preparation, the samples were run on a HiSeq 2500 machine (Illumina, US). 8 samples were run in each lane, in a paired end sequencing run, generating approximately 300 million reads across the 8 samples.

High-throughput RNA sequencing raw data were uploaded to Galaxy, an open access portal for next-generation sequencing analysis. For each condition, a N of 4 was used in each group, and three lanes of data were collected for each sample. BAM files from each of the three lanes were merged. The merged BAM files were analysed for gene expression differences using CuffDiff analysis. The CuffDiff parameters included geometric library organisation, pooled dispersion estimation and the false discovery rate was set at 0.05. Minimum alignment count – 10, multi-read correct, bias correction and CuffLinks effective length correction. Differential gene expression was calculated from fragments per kilobase per million mapped reads (FKPM) values. Differential expression was assessed across time and treatment using CuffDiff. Multiple hypothesis correction was carried out using the Benjamini-Hochberg test., Data deemed statistically significant, FDR < 0.05. Differentially expressed genes were uploaded to DAVID for gene ontology analysis.

### RNA-sequencing rhythmicity analysis

For CTL and MPL treated groups, we focused on 24h periodicity using harmonic regression on the log2 transformed signals, as previously described^48^, from gene expression data sampled every 4 hours. Each hour on the clock face refers to the total number of genes that peak at that time of day.

### Brain slice preparation

The brain was rapidly removed at either ZT0030 (30-60min following light change - inactive phase recording) or ZT1230 (30-60min following light change - active phase recording) (Suppl. Fig. 5) and placed in ice cold slicing solution containing (in mM): 52.5 NaCl, 2.5 KCl, 25 NaHCO_3_, 1.25 NaH_2_PO_4_, 5 MgCl_2_, 25 D-Glucose, 100 Sucrose, 2 CaCl_2_, 0.1 Kynurenic acid, bubbled with 95% O_2_/5% CO_2_. Para-sagittal slices (400 μm) were cut using a vibrating blade microtome (Leica VT1000 S) whilst maintained in slicing solution. The hippocampus was then isolated and placed in a holding chamber containing artificial cerebrospinal fluid (aCSF) (containing (in mM) 124 NaCl, 3 KCl, 26 NaHCO_3_, 1.4 NaH_2_PO_4_, 1 MgSO4, 10 D-Glucose, 2 CaCl_2_) where they were incubated at 32°C for 30 mins, followed by RT for at least 30 mins (for whole-cell recordings) or kept at room temperature for at least 1 hour (for field electrophysiology) before being transferred to the recording chamber.

### Extracellular electrophysiology

Hippocampal slices were placed in a submersion style recording chamber maintained at 30°C, and continuously perfused with oxygenated aCSF at a flow rate of ∼2 mL/min. Field excitatory post-synaptic potentials (fEPSPs) were evoked at 0.033 Hertz (Hz) by placing bipolar stimulating electrodes on the Schaffer collateral fibres with the recording electrode positioned in CA1 stratum radiatum. Recording electrodes were prepared by pulling borosilicate capillary tubes with a P-97 Flaming/Brown micropipette puller (Sutter Instrument Co) to a tip resistance of 2-6 MΩ and were then back-filled with aCSF. Signals were amplified using an AxoClamp 2B amplifier (Molecular Devices), digitized using a BNC-2110 (National Instruments) board, and 50/60 Hz noise eliminated by a Hum Bug (Quest Scientific). Data was acquired and analysed using WinLTP software^84^. Signals were averaged over a period of 2 minutes and a stable baseline recording of 30 minutes was acquired before LTP induction by delivery of 10 Hz stimulation for 90 seconds.

### Whole-cell electrophysiology

Whole cell recordings were taken from pyramidal neurons in the CA1 cell layer. Borosilicate pipettes (2-6 MΩ) were filled with an internal solution containing (in mM): 8 NaCl, 130 CsMeSO_4_, 10 HEPES, 0.5 EGTA, 4 MgATP, 0.3 NaGTP, 5 QX-314. Recordings were accepted for analysis with an uncompensated series resistance of <2.5 times the pipette resistance. Recordings were not corrected for series resistance due to the small current amplitudes. During mEPSC and mIPSC recordings, 100 µM D-AP5 was added to the perfused aCSF to block NMDA receptor mediated currents. mEPSCs were recorded at a membrane potential of -70 mV and for mIPSCs membrane potential was held at 0 mV. Recordings were amplified using an AxoClamp 700B (Molecular Devices) for whole-cell voltage-clamp recordings. Data was acquired using WinLTP software at a sampling rate of 10KHz, and filtered at 6 KHz, before being analyzed offline using ClampFit 9.2. mEPSCs and mIPSCs were identified when the rise time was faster than the decay time and had a peak amplitude >6 mV.

### Object location memory task

For all memory testing different rats were used at all time points and treatment. Rats were transferred to a sound-attenuating behavior room in low light (40-50 LUX on arena floor) at ZT11. Rats were left to habituate to the room for an hour, prior to starting behavior experiments. Rats were handled for 1 week in the behavior room prior to experimental start, followed by habituation to the arena without stimuli for 10 minutes daily for 5 days. Training and testing occurred in an open-top arena (50 × 90 × 100cm) made of wood. The walls inside the arena were black in colour and floor covered in sawdust. An overhead camera recorded behaviour for analysis. Exploration was scored when the rat head orientated toward the object and came within 1cm of the object. The objects were constructed from Duplo blocks, which were too heavy for the animals to displace. During training, rats were exposed to two identical objects and allowed to explore for 4 min. These objects were placed in the far side of the arena, 10cm away from the walls to allow full access around the objects. During the retention test (1h for short-term memory, 6hr for intermediate memory, or 24h for long-term memory), rats were allowed to explore for 3 min. During testing, one of the objects was moved to a new location. The position of the object was counterbalanced between rats. Total exploration time was recorded and preference for the novel item was expressed as a discrimination index.

### Activity and temperature recording

All rats were individually housed for telemetry recordings for technical reasons. Rats were implanted intraperitoneally with telemeters (PTD 4000 E-mitter, Starr Life Sciences Corp, US). Following recovery (>3 days), cages were placed upon receivers (ER-4000 receiver, Starr Life Sciences Corp, US). Locomotor activity and core body temperature data were collected every 10 minutes for 5 consecutive days. Data collected were analyzed with R CRAN package *cosinor* to measure bathyphase, acrophase, and circadian period under a 24-hour period.

### Statistical analyses

Results are presented as mean ± s.e.m. All statistical analyses were performed using GraphPad Prism (v 9.1, GraphPad software Inc., US), with parametric and non-parametric tests used where appropriate. Details of specific tests are provided in the figure legends. Statistical significance was set at *P*<0.05.

## Data availability

All raw data are available upon request to the corresponding author. All RNA sequencing data will be uploaded to the Gene Expression Omnibus (GEO) prior to publication.

